# Using single visits into integrated occupancy models to make the most of existing monitoring programs

**DOI:** 10.1101/848663

**Authors:** Valentin Lauret, Hélène Labach, Matthieu Authier, Olivier Gimenez

## Abstract

A major challenge in statistical ecology consists of integrating knowledge from different datasets to produce robust ecological indicators. To estimate species distribution, occupancy models are a flexible framework that can accommodate several datasets obtained from different sampling methods. However, repeating visits at sampling sites is a prerequisite for using standard occupancy models. Occupancy models were recently developed to analyze detection/non-detection data collected during a single visit. To date, single-visit occupancy models have never been used to integrate several different datasets. Here, we showcase an approach that combines two datasets into an integrated single-visit occupancy model. As a case study, we estimated the distribution of Bottlenose dolphins (*Tursiops truncatus*) over the North-western Mediterranean Sea by combining 24,624 km of aerial surveys and 21,464 km of at-sea monitoring. We compared the outputs of single- vs. repeated-visit occupancy models into integrated occupancy models. Integrated models allowed a better sampling coverage of species home-range, which provided a better precision for occupancy estimates than occupancy models using datasets in isolation. Overall, single- and repeated-visit integrated occupancy models produced similar inference about the distribution of bottlenose dolphins. We suggest that single-visit occupancy models open promising perspectives for the use of existing ecological datasets.

## Introduction

In large-scale ecological analysis, several parallel monitoring programs are often carried out to collect ecological data (Zipkin and Saunders 2018). Ecological monitoring programs are conducted by organizations operating across different time scales, geographic scales and funding initiatives (Lindenmayer and Likens 2010). A major challenge is integrating knowledge from different monitoring programs to produce robust ecological indicators that may be used to inform decision-making (Lindenmayer and Likens 2010, Fletcher et al. 2019). Recently, modelling tools have emerged to combine multiple data sources to estimate species distributions (Miller et al. 2019). *Integrated models* refer to the approaches that combine different data sources (Miller et al. 2019, Isaac et al. 2019). The main purpose of integrated models is to improve the accuracy of ecological indicators (Fletcher et al. 2019, Zipkin et al.2019). Species distributed over large areas could particularly benefit from integrated models because they allow for a global coverage of species occurrence by combining different data sources collected at different spatial scales (Miller et al. 2019).

To estimate species distribution in the face of uncertainties inherent to data collection, occupancy models are widely-spread statistical tools (Mackenzie et al., 2002). Occupancy models have been developed to estimate species distribution while accounting for false negatives in the observation process (Mackenzie et al. 2002). Estimating occupancy when species detection is not perfect requires performing *repeated visits* to assess the detection probability (MacKenzie 2006). However, repeating visits is sometimes unfeasible due to associated costs and logistical issues. In this context, two relevant developments of occupancy models have been recently proposed. First, integrated occupancy models combine data from different monitoring programs to improve the estimation of species distribution (Miller et al. 2019, Fletcher et al. 2019). Second, Lele et al., (2012) used occupancy models to estimate species distribution and detectability while having only one visit at the sampling site, i.e. hereafter *single-visit* occupancy models. An increasing number of studies suggest that under certain conditions, single-visit models produce robust estimates of occupancy without repeating visits at the sampling sites (Lele et al. 2012, Sólymos and Lele 2016, Peach et al. 2017). Besides, single-visit occupancy offers the possibility to work with existing datasets that did not carry out repeated visits, which is relevant to population biology and management.

In this paper, we develop an integrated approach that combines two single-visit occupancy models and demonstrate that combining several datasets into integrated single-visit occupancy models leads to accurate ecological parameter estimation. We also investigate the performance of single-visit vs. repeated-visits occupancy models. As a case study, we focused on the distribution of Bottlenose dolphins (*Tursiops truncatus*) in the North-Western Mediterranean Sea. We combined aerial surveys and at-sea monitoring into integrated occupancy models and we compared the outputs of integrated occupancy models to occupancy models using each dataset in isolation. Last, we discuss the advantages of integrated single-visit occupancy models to deal with existing ecological monitoring programs.

## Methods

### MODEL DESCRIPTION

#### Latent ecological process

Occupancy models estimate spatial distribution while accounting for imperfect species detection (Mackenzie et al. 2002). The formulation of occupancy models as state-space models allows distinguishing the latent ecological state process (i.e. species present or absent at a grid-cell) from the detection process (Royle and Kéry 2007). We denote *z*_*i*_ the latent occupancy of grid-cell *i* (*z =* 1, presence; *z =* 0, absence). We assume *z*_*i*_ is drawn from a Bernoulli distribution with *Ψ*_*i*_ the probability that the species is present at grid-cell *i*:

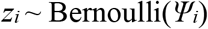

We modelled *Ψ* as a function of some environmental covariate on a logit scale, say habitat.

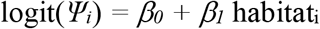

where parameters *β*_*0*_, and *β*_*1*_ are to be estimated.

### Repeated-visit observation process

In standard occupancy designs, each grid-cell is visited *J* times to estimate the detection probability. We denote *y*_*i,j*_ (*y*_*i,j*_ *=* 0, no detection ; *y*_*i,j*_ *=* 1, detection) the observations corresponding to the data collected at grid-cell *i* during visit *j* (*j =*1, ‥,J). Repeating visits at a grid-cell allows estimating species detectability, with *p*_*i,j*_ being the probability of detecting the species at visit *j* given it is present at grid-cell *i*:

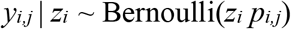

### Single-visit observation process

The difference with repeated-visit occupancy models lies in the number of sampling occasions which is *J* = 1 in single-visit occupancy models. The *j* subscript is dropped and we denoted *y*_*i*_ the observation corresponding to the data collected at site *i*. Subsequently, *p*_*i*_ is the probability of detecting the species during the single visit given it is present at site *i*:

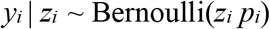

Single-visit occupancy models require certain conditions to be fulfilled for estimating detection probabilities reliably. First, different continuous covariates should be used to estimate detection and occupancy probabilities (Lele et al. 2012, Peach et al. 2017). Second, the number of detections is an important parameter that may affect the results in the case of rare or ubiquitous species (Peach et al. 2017). Third, the use of inappropriate link functions to model the detection process may lead to model misspecification and biased interpretation (Knape & Korner-Nievergelt, 2015). Despite these concerns, simulation studies have showed that situations where single-visit occupancy models fail are rare (Peach et al. 2017) and, in practice, the conditions for safe application of single-visit occupancy models are often fulfilled (Sólymos and Lele 2016). Because the number of detections is an important condition to accurately estimate single-visit occupancy parameters (Peach et al. 2017), we expect that integrated approaches will be beneficial to single-visit occupancy modelling by increasing the number of detections (true occupancy) available.

### Integrated occupancy models

We developed an integrated occupancy model using data from two independent monitoring programs, say A and B. The state process driving the latent occupancy state of site *i, z*_*i*_, remains unchanged and is drawn from a Bernoulli distribution with probabilityΨ, which is modeled as a function of environmental covariates. The observation of the targeted species at site *i* during occasion *j* may take four values with *y*_*i,j*_ = 0 for no detection, *y*_*i,j*_ = 1 for detection in dataset A, *y*_*i,j*_ = 2 for detection in dataset B, and *y*_*i,j*_ = 3 for detection in both datasets A and B. For convenience, we drop the subscripts in the notation as the formulation of the integrated observation process is identical whether we consider single-visit occupancy (i.e. *J* = 1) or repeated-visit occupancy (*J* > 1). Assuming that detection methods are independent, the observation process can be written using detection probability by the monitoring program A (*p*_*A*_) and detection probability by the monitoring program B (*p*_*B*_):

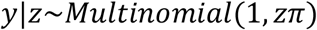

with

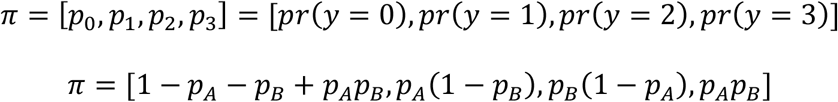

We modeled monitoring-specific detection probabilities as functions of the sampling effort of each monitoring program:

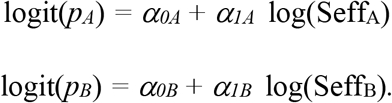

where the parameters *α*_*0A*_, *α*_*1A*_, *α*_*0B*_, and *α*_*1B*_ are to be estimated.

For example, if we assume that the detection history at site *i* is *y*_*i*_ = {2,0,1,2} over *J* = 4 sampling occasions, i.e. the species was detected by monitoring program B only at sampling occasions *j =* 1 and *j =* 4, then went undetected at *j =* 2, and was detected by monitoring program A only at *j* = 3, then for single-visit integrated occupancy we consider *y*_*i*_ *=* {3} because both monitoring programs detected the species at site *i*. We ran a simulation study comparing the performance of single-vs. repeated-visit occupancy over different scenarios affecting occupancy, and detection probabilities (Appendix S1).

## BOTTLENOSE DOLPHINS CASE STUDY

We aimed at estimating bottlenose dolphin (*Tursiops truncatus*) distribution in an area of 255,000 km^2^ covering the North-Western Mediterranean. The protected status of this species within the French seas led to the development of specific programs to monitor Mediterranean bottlenose dolphins within the implementation of the European Marine Strategy Framework Directive (2008/56/EC; MSFD). We considered two large-scale monitoring programs about bottlenose dolphins. We divided the study area in 4,356 contiguous pixel/grid-cells creating a 5’x5’ Mardsen grid (WGS 84).

We used data from at-sea surveys over 21,464 km of the French continental shelf (456 grid-cells sampled, 10.46% of the total number of grid-cells). Observers performed monitoring aboard small sailing and motor boats to locate and photo-identify bottlenose dolphins all year long between 2013 and 2015 (Labach et al. 2019). At-sea surveys detected 129 bottlenose dolphin groups located in 89 different grid-cells. At-sea surveys did not include planned repeated visits. Some sites have been visited once, and others have been visited 50 times, making this dataset a relevant candidate for single-visit model implementation.

Besides, we considered data collected during aerial line-transects covering 24,624 km of the French Exclusive Economic Zone (EEZ), targeting marine megafauna, and following a distance sampling protocol. The survey sampled 1336 grid-cells (i.e. 30.67% of the total number of grid-cells). Aerial surveys produced 130 bottlenose dolphin detections located in 87 grid-cells. Sampling effort for aerial surveys was homogeneous over the studied area with three or four replicates per line-transect between November 2011 and August 2012 (Laran et al., 2017).

An important assumption of occupancy models is that the latent ecological state of a grid-cell (the *z*_*i*_’s) remains unchanged between the repeated visits (MacKenzie 2006). When monitoring highly mobile species, such as cetaceans, the closure assumption is likely to be violated because individuals can move into and out of the sampling grid-cell. Occupied locations are used only temporarily by individuals (MacKenzie 2006; Neilson et al. 2018). Therefore, the occupancy estimator *Ψ*_*i*_ represents the probability that grid-cell *i* is *used* by the target species as opposed to the probability of occupancy (Kendall et al. 2013), and occupancy was interpreted as *space-use* by bottlenose dolphins. The detection probability now accounts for both the probability of detecting the species given that the species is available for sampling, and the probability that the species is using the grid-cell during the sampling, reflecting that the species might occupy the grid-cell but not during the sampling occasion (MacKenzie 2006). If individuals’ movement in and out of the sampling units is random, then the occupancy estimator is unbiased (Kendall et al. 2013). Occupancy and space-use refer to different scales of space selection by a species. Occupancy describes the species home range, while space-use refers to the use by individuals of the different components of the home range (Johnson 1980).

Because at-sea and aerial surveys were performed during different years, we considered them as independent. We neglected the seasonal variation in the bottlenose dolphin space-use, and we assumed that space-use remained unchanged during the monitoring period (i.e. 2011 to 2015). Concerning the ecological process, we used two environmental covariates to estimate the space-use of bottlenose dolphins: i) bathymetry, which is expected to have a positive effect on bottlenose dolphins’ occurrence (Bearzi et al. 2009, Labach et al. 2019), and ii) sea surface temperature (SST, AQUA MODIS | NASA 2019, https://neo.sci.gsfc.nasa.gov/), which is locally related to dolphins’ prey abundance and hence expected to affect local distribution of bottlenose dolphins (Bearzi et al. 2009). We extracted average SST between 2011 and 2015 value in each grid-cell, making SST a cell-specific covariate. Similarly, bathymetry had a single value per grid-cell. We checked for correlation between the two covariates and the Pearson coefficient was < 0.3. Then, we modelled *Ψ* as a function of bathymetry, SST, and the interaction between bathymetry and SST on a logit scale:

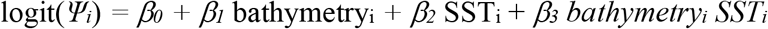

Regarding the observation process, we calculated the transect length (in km) prospected by each monitoring protocol within each grid-cell during a time period. Sampling effort was therefore a grid-cell and time-specific covariate, Seff_A_ refers to the sampling effort of the aerial monitoring program while Seff_S_ refers to the sampling effort of the at-sea monitoring program. We modeled monitoring-specific detection probabilities as functions of the relevant sampling effort:

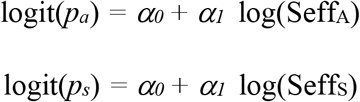

Regarding the repeated-visit occupancy models, we divided the presence-absence datasets into four sampling occasions (*J =* 4): winter (January, February, March), spring (Avril, May, June), summer (July, August, September), autumn (October, November, December). For the single-visit occupancy models, we considered the entire monitoring program in a single occasion. For example, let us assume that the detection history at site *i* is *y*_*i*_*=* {0,1,1,0} in repeated-visit occupancy, i.e. the species was detected at sampling occasions *j* = 2 and *j =* 3, and went undetected at *j* = 1, and *j =* 4, then for single-visit occupancy we have *y*_*i*_*=* {1}. In addition, the single-visit sampling effort in a grid-cell was equal to the sum of the sampling effort over the 4 sampling occasions of the repeated-visit occupancy model.

### Performances of integrated models

To assess the added value of combining aerial and at-sea datasets into integrated single-visit occupancy models, we analyzed 3 datasets: i) the aerial dataset, ii) the at-sea dataset, and iii) the two datasets together into an integrated occupancy model. For each of these datasets, we applied repeated-visit and single-visit occupancy models. Besides the case study, we also carried out a simulation study (Appendix S2).

### Bayesian implementation

We ran all models with three Markov Chain Monte Carlo chains with 100,000 iterations each in JAGS (Plummer and others 2003) called from R (R Core Team, v 3.2.5 2019) using the *r2jags* package (Su and Yajima 2015). We checked for convergence calculating the *R-hat* parameter (Gelman et al. 2013) and reported posterior means and 95% credible intervals (CI) for each regression coefficient of covariates affecting space-use probability (Fig. 1). Hereafter, we considered *effect size* of a covariate as the estimate of its regression coefficient. We discussed the effect of a covariate whenever the 95% CI of its associated parameter did not include 0. From covariates’ effect size, we calculated the predicted space-use by bottlenose dolphins (i.e. *Ψ*, Fig. 2). We reported maps of standard deviation of *Ψ* (Fig. 2B). On the maps, we displayed mean and standard deviation of *Ψ* for coastal and pelagic seas according to a 500m deep boundary that corresponds to the separation of continental shelf from the abysses. Data and codes are available on Data S1, and on GitHub at https://github.com/valentinlauret/IntegratedSingleVisitOccupancy.

**Figure 1:**
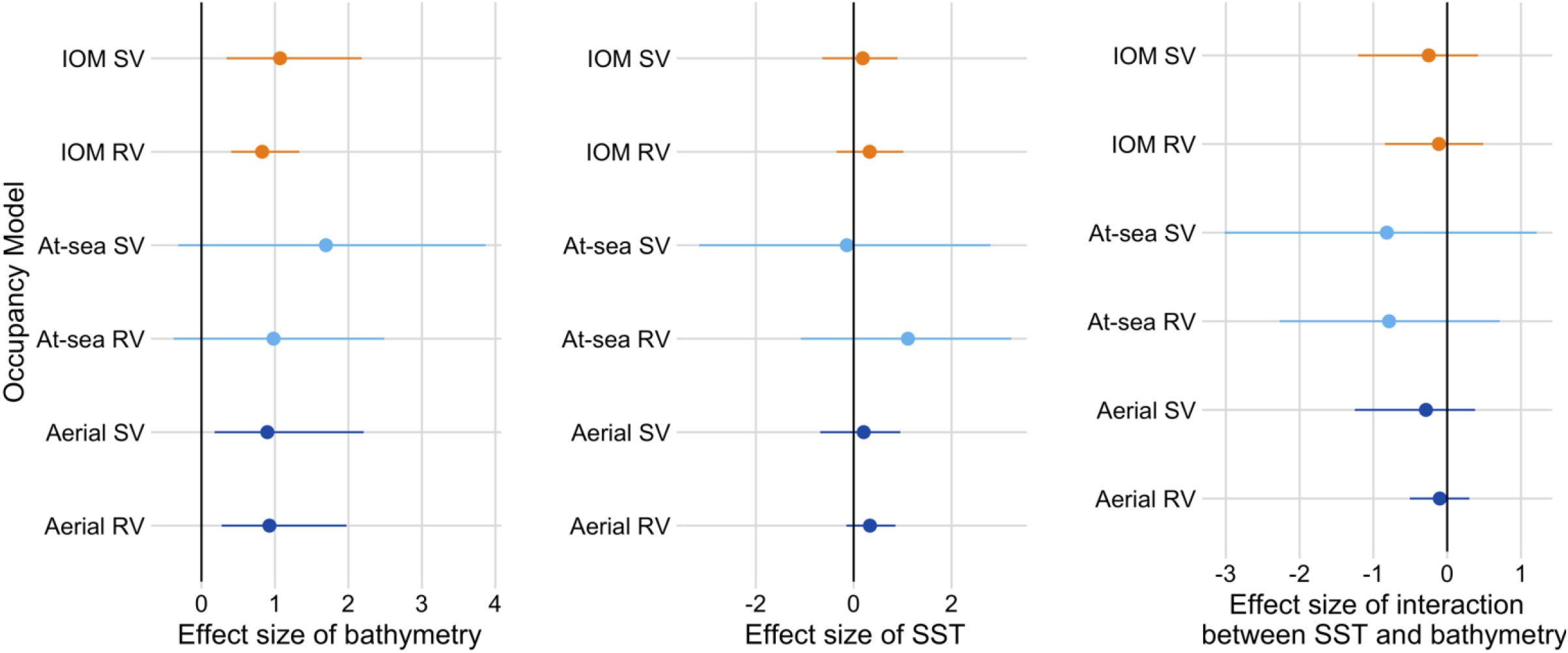
Effect size of bathymetry, sea surface temperature (SST), and interaction between SST and bathymetry on the probability *Ψ* that a site is used by Bottlenose dolphins (*Tursiops truncatus*). The posterior mean is provided with the associated 95% credible interval. “SV” refers to *single-visit* occupancy models, “RV” to *repeated visit* occupancy models, and “IOM” stands for *integrated* occupancy models, in which aerial surveys and at-sea surveys are combined. Estimates are given on the logit scale.

**Figure 2:**
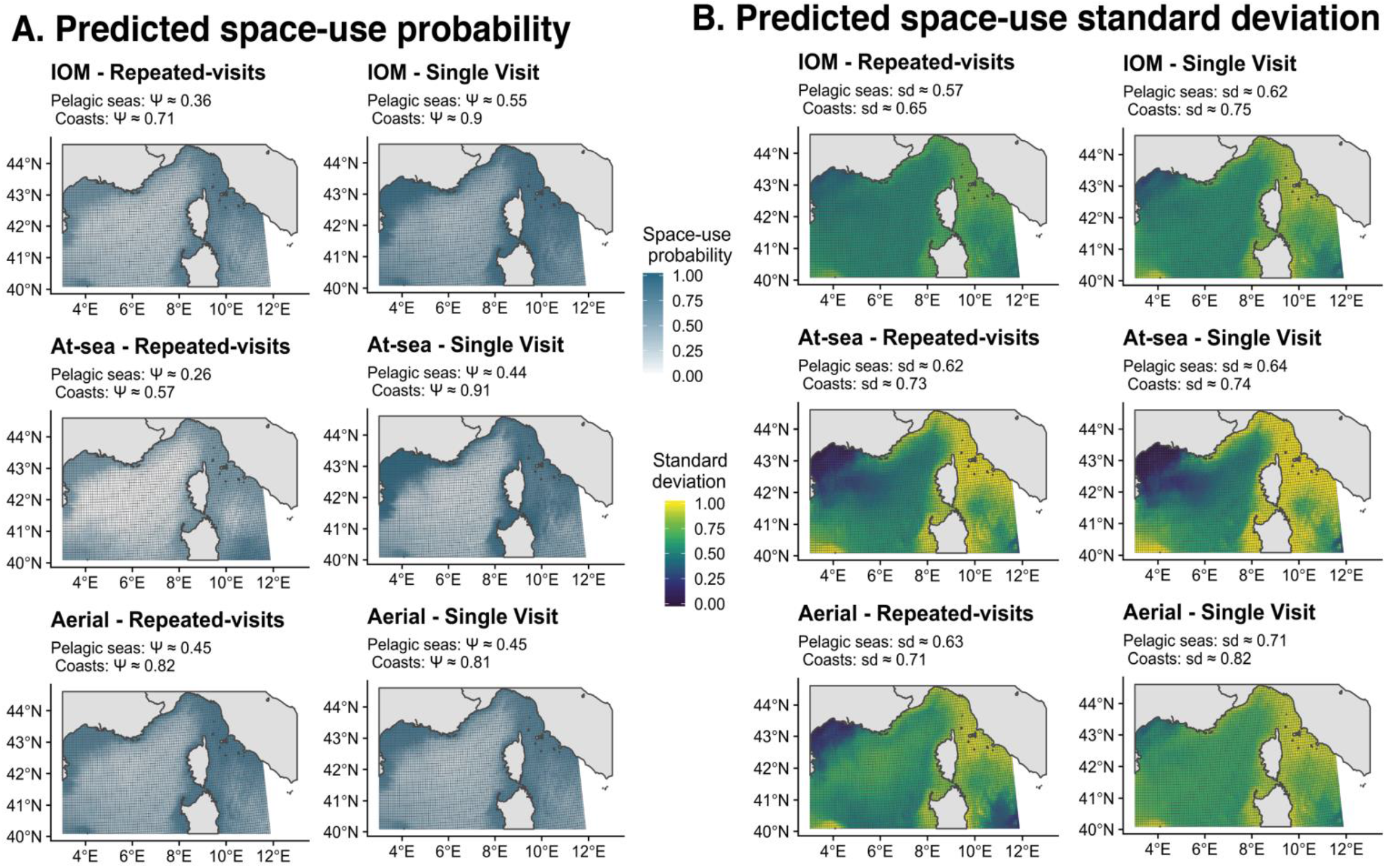
A. Probability of predicted space-use by Bottlenose dolphins (*Tursiops truncatus*) over the NW Mediterranean Sea. Using the posterior mean of covariates effect size, we estimated the probability that a grid-cell was used by bottlenose dolphins. For each occupancy model, we added the mean space-use probability (*Ψ*) for coasts (bathymetry < 500 m) and pelagic seas (bathymetry > 500 m) **B. Standard deviation of predicted space-use**. Using the posterior standard deviation of covariates effect size, we estimated the standard deviation associated with the space-use probability. For each occupancy model, we added the mean standard-deviation (sd) associated with *Ψ* for coasts (bathymetry < 500 m) and pelagic seas (bathymetry > 500 m). “IOM” stands for *integrated* occupancy models, in which aerial surveys and at-sea surveys are combined. Repeated-visit occupancy maps refer to occupancy models with 4 sampling occasions. Single-visit maps refer to occupancy models considering 1 sampling occasion.

## Results

All models produced similar predictions of space used by bottlenose dolphins (Fig. 2). The 95% CI of SST, and of the interaction between SST and bathymetry included 0 in all models (Fig. 1). The probability of space-use increased with decreasing bathymetry for all models (Fig. 1). Bathymetry ranges from altitude of 0 m to −3,488 m deep, hence a positive influence of bathymetry referred to a preference for a high seafloor (e.g. 0-200m depth). Overall, maps showed greater probabilities of space-use on the continental shelf (mean *Ψ =* 0.76 SD ± 0.17) than on the high seas (mean *Ψ =* 0.40 SD ± 0.15), although intensities of *Ψ* were different between models (Fig. 2). Bathymetry posterior means were highest for at-sea occupancy (although the 95% CI of effect size included 0), which resulted in models using only at-sea survey data predicting the highest contrast between the continental shelf and the high-seas. Bathymetry effect size was the lowest for aerial occupancy while maps from integrated occupancy models displayed moderate contrast of space-use between shelf and pelagic waters (Fig. 2). Single-visit occupancy models exhibited similar covariates estimates to those of repeated-visit occupancy models (Fig. 1). For aerial occupancy, we noticed similar space-use prediction between single- and repeated-visit (Fig. 2A). For at-sea, predicted space-use probabilities were different between single-visit and repeated-visit occupancy models (Fig. 2).

When considering the covariates’ effect size (Fig. 1), the widths of the 95% CI were not smaller for integrated occupancy than for occupancy models using datasets in isolation. However, when looking at the standard deviation of the predicted probability of space-use, integrated occupancy models had a better precision than aerial or at-sea occupancy models separately, (Fig. 2B). The use of integrated single-visit occupancy models also improved precision in predicted space-use compared to single-visit occupancy built from aerial and at-sea datasets separately (Fig.2B). Inspecting the simulation results, we found that 1) integrated occupancy models produced more precise estimates of covariates effect size than occupancy models fitted to a single dataset (Appendix S2), and 2) single-visit occupancy models produced similar results to repeated-visit occupancy models (Appendix S1).

## Discussion

### Integrated single-visit occupancy models provide reliable ecological estimates

Ecological estimates from integrated occupancy models lied within the range of the estimates obtained with each dataset separately (Fig. 1). Across all occupancy models, the effects of environmental covariates were similar and consistent with previous studies. Bottlenose dolphins were more likely to use shallower seas (Bearzi et al. 2009, Labach et al. 2019), and depth had a stronger effect than SST on the use of space by bottlenose dolphins (Torres et al. 2008). However, we found variations among models in the estimation of the probability of space-use by dolphins (Fig. 1). In particular, at-sea occupancy models predicted that dolphins make little use of the pelagic seas compared to the continental shelf, while aerial occupancy models predict more homogeneous space-use between coasts and pelagic seas. Predicted space-use was sensitive to the mean value of covariate effect size, which was similar between occupancy models. Then, variation between predicted space-use maps did not reflect the uncertainty associated with the occupancy models’ estimates. We recommend caution in interpreting predicted maps of space-use because they were mostly driven by bathymetry effect size, and did not account for precision associated with space-use prediction. To study the benefits of single-visit and integrated occupancy models to accommodate existing ecological datasets, we emphasize standard deviation maps and the credible intervals of covariates effect size (Fig. 1-2B).

Integrated occupancy models had a better precision in space-use than models using aerial or at-sea surveys separately (Fig. 2). This result was supported by our simulation study which demonstrates the better performance of integrated occupancy models at estimating covariate effect size compared to occupancy models from a single dataset (Appendix S2). Single-visit occupancy models gave similar estimates to those obtained with repeated-visit occupancy models, although repeated-visit occupancy models exhibited better precision (Fig. 1-2B), as well as in our simulations (Appendix S1).

In the bottlenose dolphins case study, we considered two existing monitoring programs that were not initially designed for occupancy modeling. In the at-sea monitoring, repeated line-transects were not implemented, nor the high depths were sampled, which made at-sea occupancy unlikely to exhibit precise estimates at our spatial extent. Besides, while aerial surveys covered a larger spatial extent than at-sea surveys, at-sea surveys exhibited a better detection rate. Detection probability was greater for at-sea surveys (p = 0.18 SD ± 0.04) than for aerial surveys (p = 0.10 SD ± 0.03). Regarding the aerial dataset, the number of occurrences was low (i.e. bottlenose dolphins were detected in 6.5% of sampled grid-cells), which might hinder the implementation of single-visit occupancy (Peach et al. 2017). Combining low-frequency occurrence data with another dataset into integrated occupancy models increases the amount of information about the ecological state process and helps mitigating the issue of low number of occurrences (the at-sea dataset had occurrences in 19.5% of sampled units).

### Ecological implications and perspectives

Overall, we illustrate that: i) Integrating datasets into occupancy models improves the precision of space-use estimates, and ii) Single-visit occupancy models can reliably accommodate the lack of repeated visits that occurs frequently. Integrated occupancy models produced more reliable estimates than occupancy models using datasets in isolation in both the bottlenose dolphin data analyzes and the simulations. Our finding on the bottlenose dolphins case study is a good illustration of the well-known benefit of combining datasets into integrated species distribution models to increased precision in ecological estimation and predicted accuracy (Fletcher et al. 2019). When the species of interest displays a wide range of occurrence (such as bottlenose dolphins), considering multiple sampling methods is effective to monitor the entire population making the most of each device (Zipkin and Saunders 2018). In particular, integrating a large volume of data, such as those that can be leveraged through citizen-science programs or with dedicated NGOs over the years can make the most of ecological monitoring programs for the furthering of many applied situations (Zipkin et al. 2019). Although repeated-visit occupancy models remain statistically more precise, there are clear benefits in using single-visit occupancy models. One advantage is the ability to relax the closure assumption, which is often incompatible with the ecological behavior of mobile species (Lele et al. 2012, Kendall et al. 2013). The closure assumption is unlikely to be valid for bottlenose dolphins over the four sampling occasions, but also for numerous monitoring programs of animal populations (Rota et al. 2009, Issaris et al. 2012, Sólymos and Lele 2016). Besides, when financial or logistical costs are limited, implementing a single-visit monitoring design could provide robust ecological estimates while explicitly accounting for imperfect species detection (Lele et al. 2012, Dénes et al. 2017).

Overall, increasing quantity and types of biodiversity data are becoming available (Isaac et al. 2019). Numerous monitoring programs do not rely on protocols implementing repeated visits like, e.g., historical monitoring programs, or citizen science programs (Tingley and Beissinger 2009, Zipkin and Saunders 2018). Then, using single-visit occupancy models helps making efficient use of available data, which is of great interest in many ecological applications (Nichols and Williams 2006, Sólymos and Lele 2016). In this context, Miller et al. (2019) encouraged further developments of methods mixing standardized and non-standardized datasets. To illustrate, we built an integrated occupancy model mixing repeated-visit occupancy models for aerial surveys and single-visit occupancy models for at-sea surveys (Appendix S3). One could also extend integrated occupancy models to more than two datasets. However, caution should be taken when integrating datasets, as combining different sources of information does not always outperform the analysis of single datasets in isolation (Simmonds et al. 2020). The flexibility of occupancy models provided a relevant framework to combine monitoring programs and to accommodate different types of data collection. Integrated and single-visit occupancy models contribute to widen the scope of possibilities. We emphasize the usefulness of both integrated and single-visit approaches to deal with existing datasets. We anticipate that their combination into integrated single-visit approaches will be of most interest for many parties in ecological research.

