## Appendix 1 for "Using single visits into integrated occupancy models to make the most of existing monitoring programs"

### Appendix S1: Integrated single-visit occupancy models

A simulation study

Valentin Lauret, Hélène Labach, Matthieu Authier, Olivier Gimenez

**Discussion** **4**

**References** **5**

---

Supporting information of the article *Using single visits into integrated occupancy models to make the most of existing monitoring programs.*

You can *download codes and results on Github*

The objective of this document is to perform a simulation study of the single-visit (SV) occupancy models and to compare their estimation to those of repeated-visits (RV) occupancy models.

SV occupancy has been developed by Lele et al. (2012). Unlike classical RV occupancy models, they support that robust occupancy estimations can be obtained from a single-visit per sampling unit. However, SV occupancy models require certain conditions to be fulfilled for estimating detection probabilities reliably. First, different continuous covariates should be used to estimate detection and occupancy probabilities (Lele et al. (2012)). Second, the number of detections is an important parameter that may affect the results in the case of rare or ubiquitous species (Peach et al. (2017)). Third, the use of inappropriate link functions to model the detection process may lead to model misspecification and biased interpretation (Knape & Korner-Nievergelt (2015)). We simulated presence-absence datasets and we aimed to compare the outputs of SV and RV occupancy models at estimating occupancy parameters. We compared SV and RV occupancy models over a large range of occupancy and detection probabilities.

### Methods

We simulated occupancy data based on a fictive covariate affecting the latent occupancy process, and 4 sampling occasions. Then, we considered two different datasets to fit occupancy models: i) a dataset with the 4 sampling occasions to fit repeated-visits occupancy model, ii) a dataset in which we considered only one sampling occasion to fit a single-visit occupancy model.

We simulated occupancy datasets with four sets of occupancy probability ($\Psi \approx 0.1$, $\Psi \approx 0.3$, $\Psi \approx 0.5$, $\Psi \approx 0.9$), and four sets of detection probabilities ($p \approx 0.1$, $p \approx 0.3$, $p \approx 0.5$, $p \approx 0.8$).

To compare model precision and bias, we calculated the relative bias (RB) and root mean square error (RMSE) of occupancy estimates over `S = 500` simulations:

- Relative bias: $RB = \frac{1}{S} \sum_1^S \frac{(\hat{\theta}_s - \theta)}{\theta}$
- Root Mean Square Error: $RMSE = \sqrt{\frac{1}{S} \sum_1^S (\hat{\theta}_s - \theta)^2}$

where $\theta_i$ is the estimate of parameter $\theta$ in the i-th simulation. We reported RB and RMSE for the regression coefficient of covariate affecting occupancy probability, and for occupancy probability itself.

#### Data simulation

##### State process

The occupancy state $z$ was drawn from a Bernoulli distribution with parameter $\psi$, $z \sim dbern(psi)$. We wrote $\psi$ as a logistic function of an environmental covariate `cov`:

$$\text{logit}(\psi) = \alpha_0 + \alpha_1 \text{ cov}$$

where $\alpha_0$ and $\alpha_1$ are unknown parameters that need to be estimated.

We considered 4 sets of values for the alpha's:

- $\alpha_0 = -1.9$ and $\alpha_1 = 0.2$ that led to $\psi$ approx. equal to 0.1
- $\alpha_0 = -0.5$ and $\alpha_1 = 0.2$ that led to $\psi$ approx. equal to 0.3
- $\alpha_0 = 0.5$ and $\alpha_1 = 0.2$ that led to $\psi$ approx. equal to 0.5
- $\alpha_0 = 2.5$ and $\alpha_1 = 0.15$ that led to $\psi$ approx. equal to 0.9

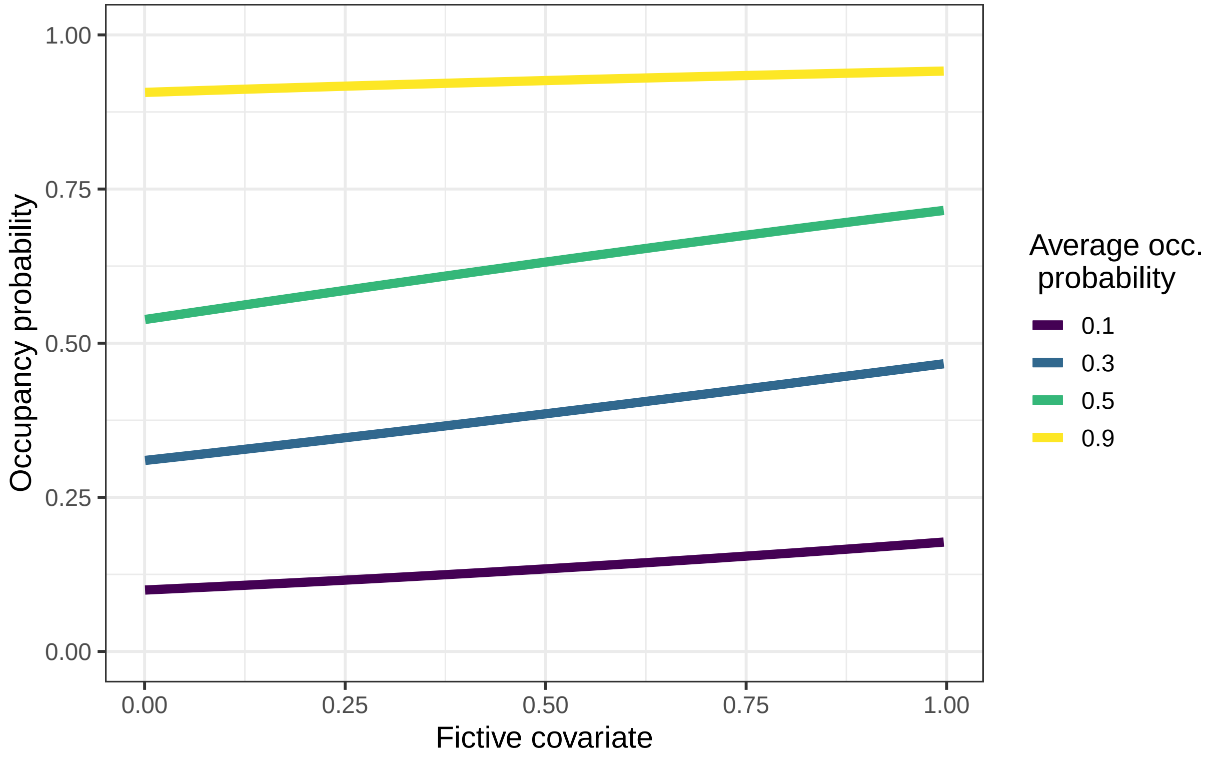

*Figure 1: Occupancy probability as a function of a fictive covariate*

###### Observation process

Observations are drawn from a Bernoulli distribution with parameter $p$. We wrote $p$ as a logistic function of a sampling effort covariate seff:

$$\text{logit}(p) = \beta_0 + \beta_1 \text{ seff}$$

where $\beta_0$ and $\beta_1$ are unknown parameters that need to be estimated.

We considered 4 sets of values for the beta's:

- $\beta_0 = -1.5$ and $\beta_1 = 0.26$ that led to $p$ approx. equal to 0.15
- $\beta_0 = -0.6$ and $\beta_1 = 0.25$ that led to $p$ approx. equal to 0.35
- $\beta_0 = 0.3$ and $\beta_1 = 0.26$ that led to $p$ approx. equal to 0.5
- $\beta_0 = 1.8$ and $\beta_1 = 0.3$ that led to $p$ approx. equal to 0.8

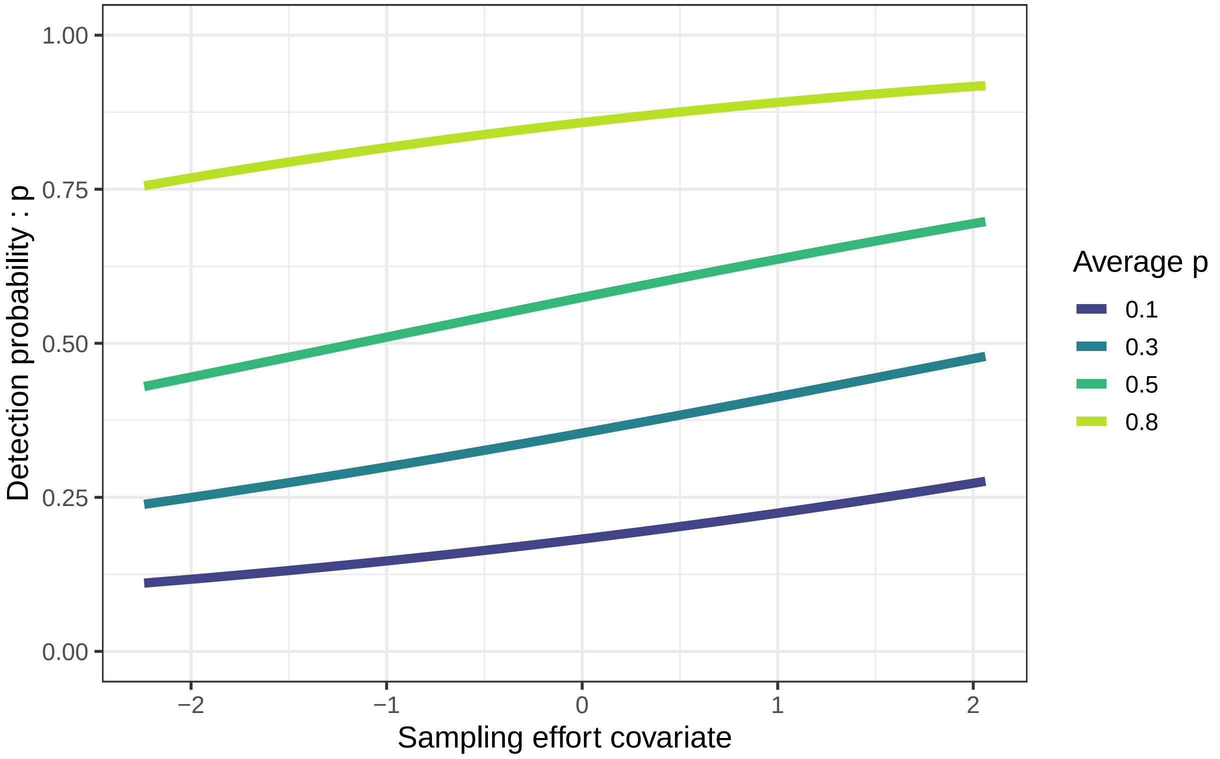

*Figure 2: Detection probability as a function of a fictive sampling effort*

###### Study area

We simulated $z$ and $y$ for study areas of 100 sites.

#### Models

For each combination of $\psi$ and $p$ we fitted 2 occupancy models:

- Repeated-visit occupancy model (RV)
- Single-visit occupancy model (SV)

We have 32 ecological situations depending on $\psi$ and $p$.

For each scenario, we ran 500 simulations and fitted the 2 occupancy models.

### Results

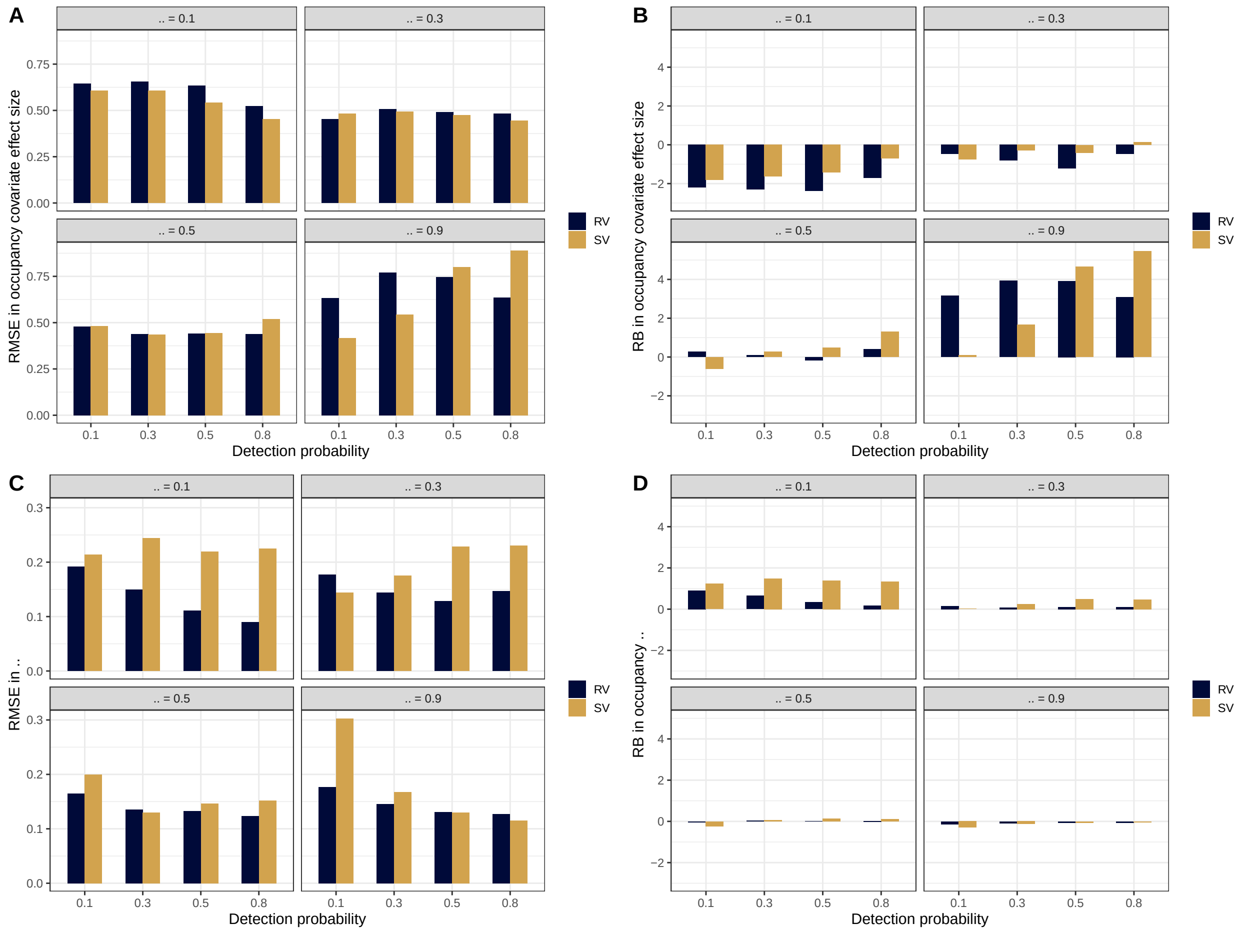

*Figure 3: Root-mean square error (RMSE) and Relative Bias (RB) of repeated- and single-visit occupancy models based on simulated data*

#### Covariate effect size

Regarding the covariate effect size on $\psi$ (Fig.3A-B), we found close RMSE and RB between repeated- and single-visit occupancy. In the case, of high occupancy ($\psi = 0.9$), differences in RB are greater although they are close in precision. Overall, the results were similar whatever the occupancy models we considered.

#### Occupancy prediction

Predicted occupancy probability between single- and repeated-visits models are closed to each other in Fig. 3C-D. However, precision and bias are greater for low occupancy $\psi = 0.1$, which is consistent with Peach et al. (2017) findings. Same results when detection probability is low and when $\psi$ is high. Although, note that RMSE plotting scale is smaller in Fig. 3C than in Fig. 3A.

### Discussion

Our simulations study showed that single-visit and repeated-visit occupancy models had similar results in the estimation of covariate effect size for ocucpancy. We explored simple logistic regressions to describe models paramters. To be consistent with our manuscript, we reported performances of covariates effect size and predicted occupancy. To have a complete understanding of models performances, one might want to look at wider range of relationships, and observe all models parameters (e.g. intercept, sampling effort effect size on detection probability).

Overall, our simulation results support that single-visit occupancy models can be used to obtain reliable estimates of occupancy.
