## Appendix 2 for "Using single visits into integrated occupancy models to make the most of existing monitoring programs"

### Appendix S2: Integrated single-visit occupancy models

A simulation study

Valentin Lauret, Hélène Labach, Matthieu Authier, Olivier Gimenez

#### Contents

**Methods** **1**
Data simulation . . . . . . . . . . . . . . . . . . . . . . . . . . . . . . . . . . . . . . . 2
Models . . . . . . . . . . . . . . . . . . . . . . . . . . . . . . . . . . . . . . . . . . . 2

**Results** **3**

**Discussion** **3**

**References** **4**

---

Supporting information of the article *Using single visits into integrated occupancy models to make the most of existing monitoring programs.*

You can *download codes and results on Github*

The objective of this document is to perform a simulation study to assess the performance of integrated occupancy models. We explored whether single-visit (SV) or repeated-visits (RV) occupancy models benefit from data integration, and we explored the effect of number of detections to estimate the SV occupancy model parameters.

### Methods

We simulated occupancy data based on a fictive covariate affecting the latent occupancy process, and 4 sampling occasions. Then, we considered three different datasets to fit occupancy models: i) a dataset with the 4 sampling occasions to fit RV occupancy model, ii) a dataset in which we pooled the 4 occasions to fit a SV occupancy model dealing with the entire dataset simulated, and iii) a dataset using only the first sampling occasion simulated to fit a single-visit occupancy model with a reduced number of detections compared to ii).

We simulated occupancy datasets with two sets of value for occupancy probability ($\Psi \approx 0.1$, and $\Psi \approx 0.3$), and two sets of detection probabilities: $p \approx 0.1$, $p \approx 0.5$. Finally, we analysed the datasets of detection probability $p \approx 0.1$, and $p \approx 0.5$ jointly into integrated occupancy models.

To compare model precision and bias, we calculated the relative bias (RB) and root mean square error (RMSE) of occupancy estimates over `S = 500` simulations:

- Relative bias: $RB = \frac{1}{S} \sum_1^S \frac{(\hat{\theta}_s - \theta)}{\theta}$
- Root Mean Square Error: $RMSE = \sqrt{\frac{1}{S} \sum_1^S (\hat{\theta}_s - \theta)^2}$

where $\theta_i$ is the estimate of parameter $\theta$ in the i-th simulation. We reported RB and RMSE for the regression coefficient of covariate affecting occupancy probability, and for occupancy probability itself.

#### Data simulation

##### State process

The occupancy state $z$ was drawn from a Bernoulli distribution with parameter $\psi$, $z \sim dbern(psi)$. We wrote $\psi$ as a logistic function of an environmental covariate `cov`:

$$\text{logit}(\psi) = \alpha_0 + \alpha_1 \text{ cov}$$

where $\alpha_0$ and $\alpha_1$ are unknown parameters that need to be estimated.

We considered 2 sets of values for the alpha's:

- $\alpha_0 = -1.9$ and $\alpha_1 = 0.2$ that led to $\psi$ approx. equal to 0.1
- $\alpha_0 = -0.5$ and $\alpha_1 = 0.2$ that led to $\psi$ approx. equal to 0.3

##### Observation processes

Observations are drawn from a Bernoulli distribution with parameter $p$. We wrote $p$ as a logistic function of a sampling effort covariate seff:

$$\text{logit}(p) = \beta_0 + \beta_1 \text{ seff}$$

where $\beta_0$ and $\beta_1$ are unknown parameters that need to be estimated.

We considered 2 sets of values for the beta's:

- $\beta_0 = -1.5$ and $\beta_1 = 0.26$ that led to $p$ approx. equal to 0.1
- $\beta_0 = 0.3$ and $\beta_1 = 0.26$ that led to $p$ approx. equal to 0.5

We simulated 3 different datasets to fit occupancy models:

- RV occupancy dataset considering four sampling occasions (`J=4`), hereafter 'RV'
- SV occupancy dataset considering the same four sampling occasions as a single occasion (`J=1`), hereafter 'SV4'
- SV occupancy dataset considering only the first sampling occasion (`J=1`), herafter 'SV1'

#### Models

For each value of $\psi$ ($\Psi \approx 0.1$, and $\Psi \approx 0.3$), we built RV, SV4, and SV1 datasets with:

- low detection probability simulations, $p \approx 0.1$
- high detection probability simumations, $p \approx 0.5$
- both datasets combined in an integrated dataset.

Overall, we obtained occupancy models to 18 datasets.

*Note that for the integrated occupancy, we combined datasets generated from the same z ecological states with two different detection probabilities, i.e. we combined one monitoring with 0.5 detection probability (i.e. 'low p'), and one monitoring with 0.8 detection probability (i.e. 'high p').*

### Results

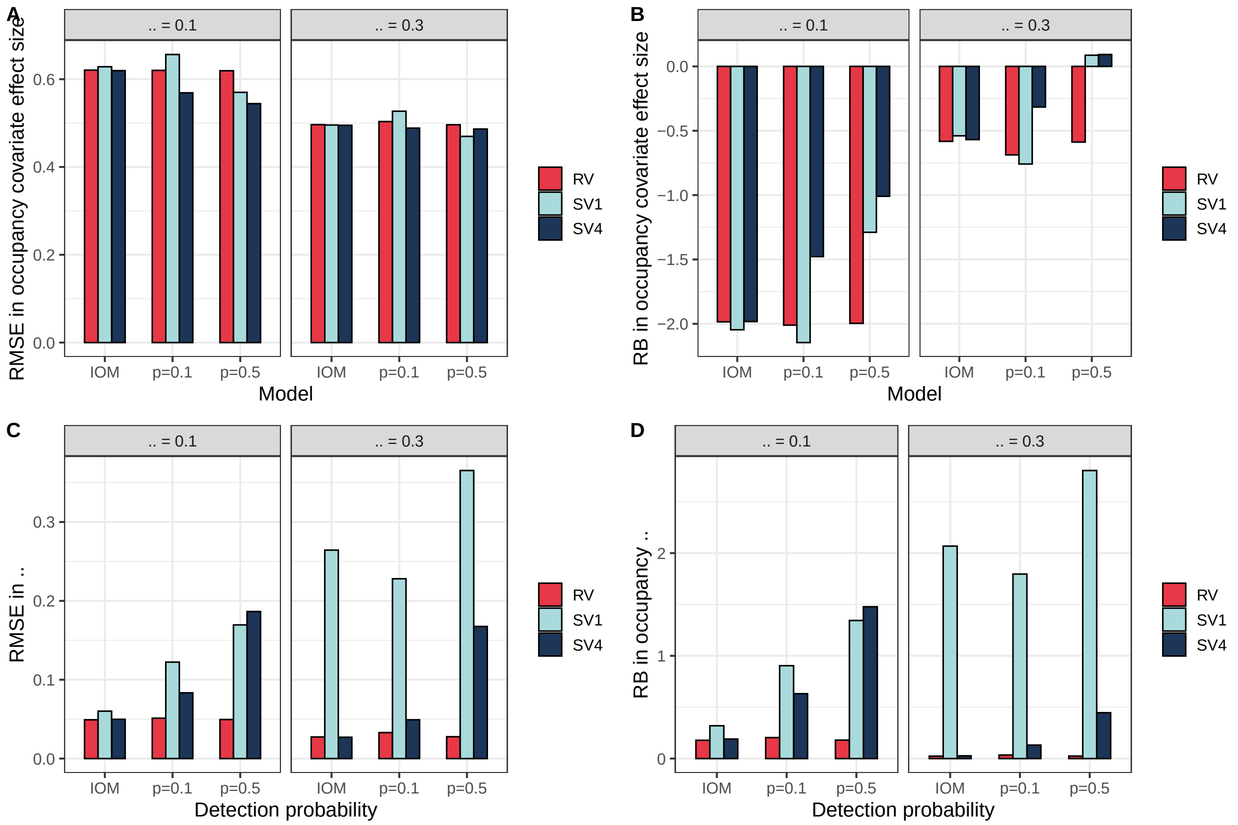


*Figure 1: Root-mean square error (RMSE) and Relative Bias (RB) of occupancy models based on simulated data. IOM stands for "Integrated Occupancy Models"*

Regarding the covariate effect size on occupancy probability (Fig. 1A-B), the RMSE values were similar whatever the occupancy model we considered. Overall, we found a better precision and bias for higher occupancy ( = 0.3). When reconstructing the occupancy probability  (Fig. 1C-D), the results between pooled single-visit and repeated-visits were similar to each other. In contrast, single-visit occupancy models using only the 1st sampling occasion exhibited a greater RMSE and RB. For pooled single-visit and repeated-visit occupancy models, integrated occupancy models exhibited better precision and bias than occupancy models using the dataset in isolation.

### Discussion

Our simulations showed that integrated occupancy models improved precision and bias of occupancy probability compared to models using the datasets in isolation. The lower performance of SV occupancy models using only the first sampling occasion is due to the limited number of detections in the dataset. We suggest that care should be taken when considering SV occupancy for datasets that arise from reduced sampling effort coupled with low detection probability, which can produce small numbers of detections which, in turn, leads to degraded performance of single-visit occupancy models. To overcome this issue, reducing the amount of sampling effort per grid-cell could be balanced by an increase in the number of sampled sites not to decrease precision in occupancy estimates. Results from both Peach et al. (2017) and our simulation study comparing single- and repeated-visit pointed out the limited performances of single-visit occupancy models in the case of low occupancy (Supporting information 1), and when the number of detections is limited.

Overall, our simulations underline that integrated single-visit occupancy models can be used to obtain reliable estimates of occupancy.
