## Appendix 3 for "Using single visits into integrated occupancy models to make the most of existing monitoring programs"

### Appendix S3: Integrated single-visit occupancy models

A simulation study

Valentin Lauret, Hélène Labach, Matthieu Authier, Olivier Gimenez

**Results** **3**

**Discussion** **3**

**References** **5**

---

Supporting information of the article *Using single visits into integrated occupancy models to make the most of existing monitoring programs.*

You can *download codes and results on Github*

The objective of this document is to explore how to build a "mixed" integrated occupancy model analyzing a repeated-visit dataset along with a single-visit dataset.

### Methods

Based on the bottlenose dolphin (*Tursiops truncatus*) case study of the paper *Using single visits into integrated occupancy models to make the most of existing monitoring programs*, we built a 'mixed' integrated occupancy models considering a repeated-vists (RV) observation process for the aerial-dataset, and a single-visit (SV) observation process for the at-sea dataset.

#### State process

The occupancy state $z$ was drawn from a Bernoulli distribution with parameter $\psi$, $z \sim \text{Bernoulli}(\psi)$. We wrote $\psi$ as a logistic function of two environmental covariates bathy (bathymetry), and SST (Sea Surface Temperature):

$$\text{logit}(\psi) = \alpha_0 + \alpha_1 \text{ bathy} + +\alpha_2 \text{ SST}$$

where $\alpha_0$, $\alpha_1$, and $\alpha_2$ are unknown parameters that need to be estimated.

#### Observation process

The observations are drawn from a Bernoulli distribution with parameter $p$. We wrote $p$ as a logistic function of a sampling effort covariate seff:

$$\text{logit}(p) = \beta_0 + \beta_1 \text{ seff}$$

where $\beta_0$ and $\beta_1$ are unknown parameters that need to be estimated.

#### Mixed observation process

In this 'mixed' integrated occupancy model, the state process remains enchanged. However, there are 2 separated observation processes, while we analyzed jointly the detections into the same observation process in the 'classical' integrated model that considered the same number of sampling occasions (`J`). The two observation processes informed the latent ecological layer.

We separated the RV aerial detections $ya$ (with 4 sampling occasions, `J=4`), from the SV at-sea detections $ys$ (i.e. 1 sampling occasion, `J=1`).

Both $ya$ and $ys$ are binary datasets (O/1) modeled as draws in Bernoulli distributions, with associated detection probabilities $pa$, and $ps$. For grid-cell $i$, during sampling occasion $j$:

- $ya_{i,j} \sim \text{Bernoulli}(z_i \; pa_{i,j})$
- $ys_i \sim \text{Bernoulli}(z_i \; ps_i)$

#### BUGS model

Hereafter, you would find the JAGS formulation of this occupancy model.

```
# Specify model in BUGS language
sink("mixed_iom.jags")
cat("
    model {

   # priors

    alpha.psi ~ dnorm(0,0.444) # occupancy intercept
    alpha.pa  ~ dnorm(0,0.444) # detection aerial intercept
    alpha.ps  ~ dnorm(0,0.444) # detection at-sea intercept

    beta.sst  ~ dnorm(0,0.444) # slope sst effect
    beta.bathy~ dnorm(0,0.444)  # slope bathy effect
    beta.eff.a~ dnorm(0,0.444)  # slope aerial survey effort effect
    beta.eff.s~ dnorm(0,0.444)  # slope at-sea survey effort effect
    beta.occ2 ~ dnorm(0,0.444) # occasion effect
    beta.occ3 ~ dnorm(0,0.444) # occasion effect
    beta.occ4 ~ dnorm(0,0.444) # occasion effect

    # State process

   for (i in 1:nsite){
      z[i] ~ dbern(psi[i])

     logit(psi[i]) <- lpsi[i]
```

```
    lpsi[i] <- alpha.psi + beta.sst * SST[i] + beta.bathy * BATHY[i]

  }  # i

     # Detection process

    # At-sea monitoring
    for(i in 1:nsite){

     mu.p_s[i] <- z[i] * p_s[i]

    logit(p_s[i]) <- alpha.ps + beta.eff.s*eff.s[i]

   y_s[i] ~ dbern(mu.p_s[i])

    } #i

    # Aerial monitoring
    for(i in 1:nsite){
      for (j in 1:nrep){

            mu.p_a[i,j] <- z[i] * p_a[i,j]

            logit(p_a[i,j]) <- lp_a[i,j]

            lp_a[i,j] <- alpha.p_a + beta.eff.a * eff.a[i,j] + beta.occ2 * equals(j,2) + beta.occ3 * equ

            y_a[i,j] ~ dbern(mu.p_a[i,j])

    } #j

  } #i

    }#fin du modele
    ", fill = TRUE)

sink()
```

### Results

Hereafter, we displayed the effect size of the environmental covariate on the estimated occupancy probability ($\psi$).

The Mixed IOM model displayed similar estimates of effect size to estimates obtained from other occupancy models presented in the manuscript. We considered that the Mixed - IOM model had a better performance than the IOM SV model. Mixed model exhibited a better precision of the covariate effect size on $\psi$ than SV integrated model on (Fig. 1), but precision was equivalent to RV integrated model.

### Discussion

This extension of SV and RV integrated occupancy modles highlights the flexibility of occupancy model to fit with the sampling designs of existing datasets. However, the separated formulation of the detection process into 2 Bernoulli draws is relevant only if the monitoring programs are independant. Dependance between

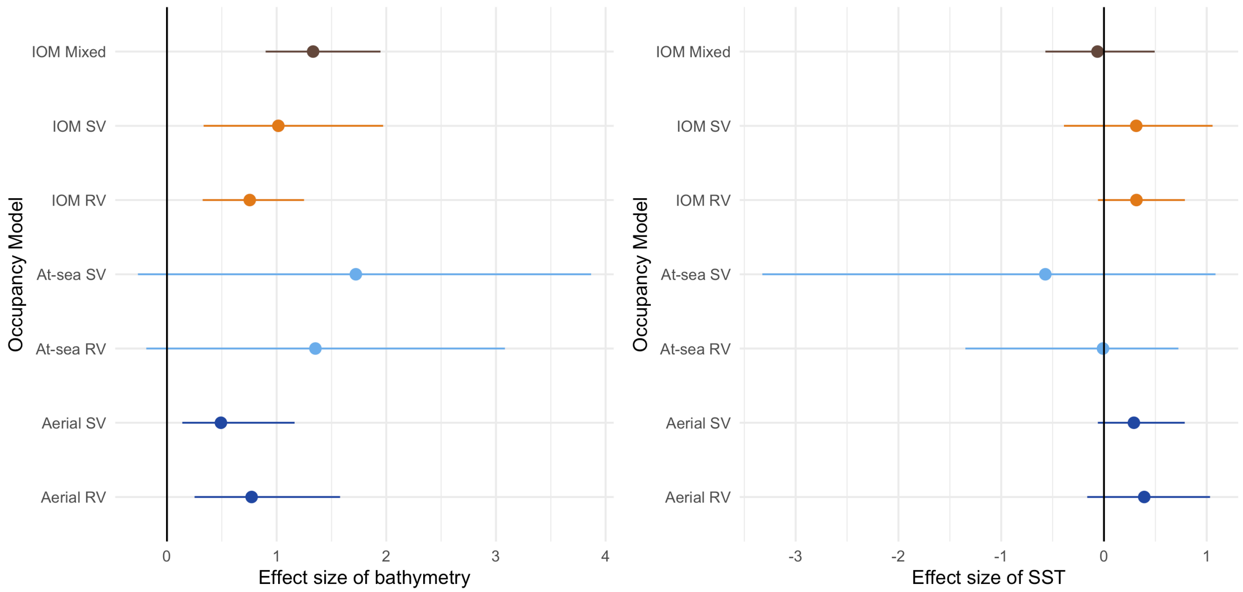

Figure 1: Figure 1 :Effect size of sea surface temperature (SST) and bathymetry on the space-use probability. Posterior mean is given with the associated 95% credible interval. Estimates are given on the logit scale. "SV" refers to single-visit occupancy models. "RV" refers to repeated-visit occupancy models. "IOM" stands for integrated occupancy models, in which aerial surveys and at-sea surveys are combined. "IOM - Mixed" refers to the 'mixed' model described above in this document.

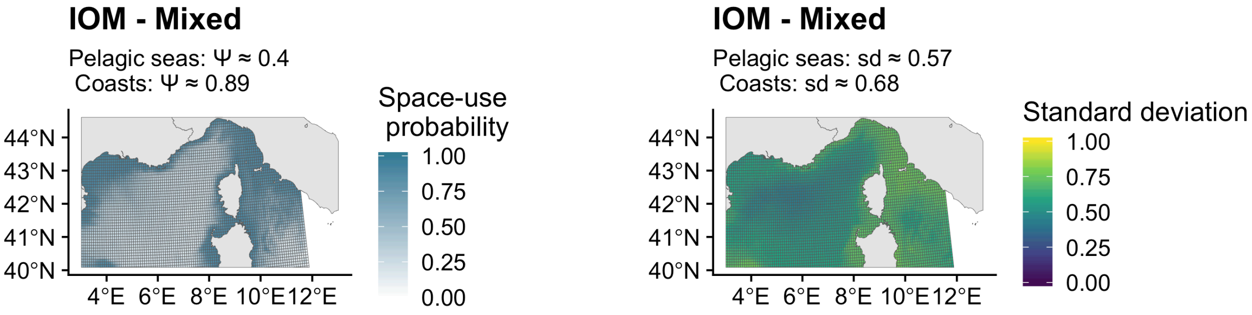

Figure 2: Figure 2: Map of predicted space use and associated standard deviation for the mixed model

monitoring devies requires to model explicitly the covariation between detection probabilities Clare et al. (2017).

Miller et al. (2019) encouraged further developments of methods mixing standardized and non-standardized datasets. We support that occupancy models provide a relevant framework to integrate monitoring programs and to accommodate different types of data collection. Integrated and single-visit occupancy models contribute to widen the scope of possibilities.
